# Beyond thermodynamic constraints: Evolutionary sampling generates realistic protein sequence variation

**DOI:** 10.1101/180331

**Authors:** Qian Jiang, Ashley I. Teufel, Eleisha L. Jackson, Claus O. Wilke

## Abstract

Biological evolution generates a surprising amount of site-specific variability in protein sequences. Yet attempts at modeling this process have been only moderately successful, and current models based on protein structural metrics explain, at best, 60% of the observed variation. Surprisingly, simple measures of protein structure, such as solvent accessibility, are often better predictors of site-specific variability than more complex models employing all-atom energy functions and detailed structural modeling. We suggest here that these more complex models perform poorly because they lack consideration of the evolutionary process that is in part captured by the simpler metrics. We compare protein sequences that are computationally designed to sequences that are computationally evolved using the same protein-design energy function and to homologous natural sequences. We find that by a wide variety of metrics, evolved sequences are much more similar to natural sequences than are designed sequences. In particular, designed sequences are too conserved on the protein surface relative to natural sequences whereas evolved sequences are not. Our results suggest that evolutionary simulation produces a realistic sampling of sequence space. By contrast, protein design—at least as currently implemented—does not. Existing energy functions seem to be sufficiently accurate to correctly describe the key thermodynamic constraints acting on protein sequences, but they need to be paired with realistic sampling schemes to generate realistic sequence alignments.

## Introduction

Alignments of protein sequences can display large amounts of position-specific variability. This variability exists because selective pressures for proteins to fold and function are not experienced uniformly across a protein’s sequence. The relative variability of a position in a sequence is often associated with where that position is located in the protein’s structure (Thorne 2007; Liberles *et al.* 2012; Arenas *et al.* 2015; Echave *et al.* 2016). For example, patterns of sequence variation differ between the core and surface of proteins. The core tends to evolve more slowly and be more conserved than sites on the surface (Kimura and Ohta 1974; Overington *et al.* 1992; Mirny and Shakhnovich 1999; Franzosa and Xia 2009; Ramsey *et al.* 2011; Tóth-Petróczy and Tawfik 2011). These core sites also tend to have more hydrophobic residues while sites on the surface tend to have more polar residues (Jones and Thornton 1996; Bastolla *et al.* 2007; Jackson *et al.* 2013). In general, sites that are conserved are assumed to be more important for protein structure, stability, and ultimately function, even though most sites in a protein are not directly involved in function.

While a significant body of literature exists examining how protein structure and thermodynamic constraints interact with evolutionary processes to shape site-specific variability, current models explain, at best, 60% of variability (Echave *et al.* 2016). Existing models span from ones using very simple metrics, such as relative solvent accessibility (RSA) or weighted contact number (WCN), to ones using complicated all-atom energy functions and protein design. One surprising result from these efforts is that the simple models tend to perform as well as or better than the complicated models. In particular, protein design has worked particularly poorly in predicting site variability observed in natural sequence alignments (Jackson *et al.* 2013; Shahmoradi *et al.* 2014; Jackson *et al.* 2016).

Why the more complex models fail to explain observed sequence variation is unknown. All-atom stability models should account for the same properties reflected in RSA and WCN, as well as an array of other features. For example, the mean interaction energy of a site with the rest of the protein is a function of a site’s contacts, reflected in WCN (Halle 2002). And thermodynamic stability is related to RSA through the energetic cost of burying a side chain in the protein core (Zhou and Zhou 2004). At the same time, all-atom models remain limited in important aspects. They generally do not accurately estimate the entropic contribution of the unfolded protein ensemble, and they tend to become increasingly inaccurate as more changes are introduced into the original protein sequence.

In addition to any biophysical limitations exhibited by all-atom models, there is another critical aspect by which protein design methods deviate from the process of protein evolution as it plays out in living organisms: Under evolution, mutations are introduced one by one and either fix or are lost to drift. As a consequence, a mutation that is deleterious in the current genetic background will rarely fix, even if it were acceptable in other genetic backgrounds. Protein design, by contrast, allows the replacement of multiple amino acids at once, and thus theoretically can create sequence configurations that are thermodynamically allowed but not easily accessible by evolution (Huang *et al.* 2016). In fact, a number of recent studies have begun to examine how the state of the background sequence affects a protein’s ability to accumulate substitutions (Pollock *et al.* 2012; Shah *et al.* 2015; Goldstein and Pollock 2016; McCandlish *et al.* 2016). These studies suggest that the order in which changes to a sequence occur can affect a position’s propensity to tolerate further mutations. Therefore, a protein’s evolutionary history may influence how mutational space is sampled, and the order in which states were sampled and the length of time any specific state has been resident may also influence sampling behavior. Hence, the effect the evolutionary process has on shaping site-specific variability could be considerable.

Here we explore the influence of evolutionary history in the generation of site-specific variability. By comparing protein sequences produced under a model that combines evolutionary history and stability constraints to sequences produced by a model that considers stability constraints alone, we can begin to tease apart the relationship between evolutionary history, protein stability, and sequence variability. To generate sequences in the absence of evolutionary history, we employ protein design as implemented in the RosettaDesign suite (Kuhlman *et al.* 2003). In this method, all of the residues are replaced simultaneously. To add evolutionary history, we use the same Rosetta force field but introduce substitutions sequentially, in a process that mimics evolution under natural selection (Teufel and Wilke 2017). We then compare the sequences produced by each of these methods to one another and to natural sequences. We find that proteins generated by simulating evolution display similar site-specific variability to natural proteins whereas designed sequences do not.

## Methods

### Protein structures and alignments

We analyzed a set of 38 protein with available structures from *Saccharomyces cerevisiae*. This dataset had previously been studied in the context of protein design by Jackson *et al.* (2013), and it had originally been assembled by Ramsey *et al.* (2011). For each structure, Ramsey *et al.* (2011) had also assembled alignments of homologous sequences containing at least 50 sequences each. Table S1 lists the protein data bank identifier (PDB ID) for each of the template structures and the number of homologous sequences available in the respective alignment.

### Generation of protein sequences

For each of the 38 protein structures, we computationally generated alignments of 500 variant sequences both via protein design and via protein evolution. In all cases, we first minimized the structures with Rosetta (Leaver-Fay *et al.* 2011). This minimization ensures that the energy difference observed upon subsequent mutation can be attributed to the effect of the mutation and is not simply caused by improved amino-acid side-chain packing. Unless explicitly noted otherwise, we initialized both the protein design and the protein evolution simulations with the exact structures and amino-acid sequences corresponding to the PDB and chain identifiers listed in Table S1.

For protein design, we used the fixed-backbone method implemented in RosettaDesign (Kuhlman *et al.* 2003). This method only allows for movement of the side chains, while the backbone is held fixed. The following command was used:

~~~
./fixbb.linuxgccrelease-database rosetta_database
        -s input.pdb-resfile ALLAA.res-ex1-ex2
        -extrachi_cutoff 0-nstruct 1
        -overwrite-minimize_sidechains
        -linmem_ig 10
~~~

For evolutionary simulations, we used an accelerated origin– fixation algorithm (Kachroo *et al.* 2015; Teufel and Wilke 2017). In an origin–fixation model, mutations are sequentially introduced and either accepted or rejected based on their effect on fitness. Under the accelerated version of this model, beneficial mutations are always fixed while deleterious mutations are exponentially suppressed. We have previously shown that this accelerated algorithm produces the same steady-state distribution of genotypes visited as the regular, non-accelerated algorithm (Teufel and Wilke 2017).

To calculate stabilities for proteins during simulated evolution, we used the `get_fa_scorefxn` function in the PyRosetta (Chaudhury *et al.* 2010), and interpreted its result as the stability Δ*G* of each proposed mutation. To convert protein stability into fitness, we use a soft-threshold model (Chen and Shakhnovich 2009; Wylie and Shakhnovich 2011; Serohijos *et al.* 2012), where the fitness *ƒ*_*i*_ of a protein *i* with stability Δ*G*_*i*_ is given by

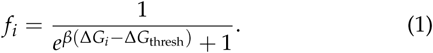

Here, *β* is the inverse temperature, Δ*G*_*i*_ is the stability of the mutant protein, and Δ*G*_thresh_ is the stability value at which fitness has declined to 50% of its maximum. For the majority of our simulations, we set Δ*G*_thresh_ to the average score obtained for the proteins designed on the same template structure. This was done to insure that proteins generated by the evolutionary model have similar stabilities as those produced by the design method. In an additional set of simulations, we set Δ*G*_thresh_ to the maximum score (i.e., corresponding to the least stable protein) obtained for the designed proteins.

To calculate the probability of fixation of a new mutation, we first log-transform fitness,

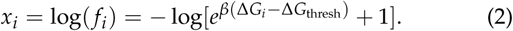

We then calculate the probability of fixation of a new mutation *j* in a background population of genotypes *i* as

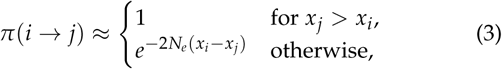

where *N*_*e*_ is the effective population size. In all simulations, we set *N*_*e*_ = 100 and *β* = 1.

We used a uniform mutation model throughout, i.e., every amino acid was equally likely to mutate into every other amino acid. However, we disallowed mutations to or from cystenines, because cysteine disulfide bonds are not properly handled by the Rosetta energy function. We ran each simulation until 5000 substitutions had occurred. Simulations were run for a fixed number of substitutions to insure similar amounts of divergence from the each of the starting templates, allowing for a fair comparison to the designed sequences. However, we note that this choice is equivalent to selecting a random substitution rather than selecting a sequence at a random time point, and it enriches for genotypes with high substitution rates.

### Data analysis

#### Site-specific variability and amino acid distributions

We separately aligned the 500 resulting sequences produced by each method for each of the 38 structures. To quantify the variability of sites in these alignments, we calculated the site entropy

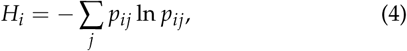

where *p*_*ij*_ is the frequency of amino acid *j* at column *i* in the alignment. Exponentiating *H*_*i*_, we obtain the effective number of amino acids,

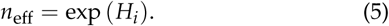

This number falls between 1 and 20 and can be interpreted as the number of different amino-acid types present at a given site.

To compare an amino-acid distribution to a reference distribution (e.g., to compare the amino-acid distribution of designed sequences to that of natural sequences), we used the Kullback-Leibler (KL) divergence, defined as

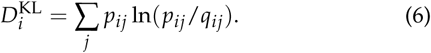

Here *q*_*ij*_ is the frequency of amino acid *j* in column *i* of the sequence alignment to compare, and *p*_*ij*_ is the relative frequency in the reference alignment. If any *q*_*ij*_ or *p*_*ij*_ were zero, we added 1/20 to each amino acid count before calculating the frequencies. To compare natural alignments to themselves, we randomly split each alignment into two equal-sized sets of sequences and then calculated the KL divergence of the first half against the second.

#### Residue buriedness

To estimate a residue’s buridness or exposure we calculated its relative solvent accessibility (RSA). RSA ranges from from 0 for completely buried residues to 1 for completely exposed ones (Tien *et al.* 2013). We first calculated the Accessible Surface Area (ASA) of each residue in each structure, using the software DSSP (Kabsch and Sander 1983). ASA indicates the surface area of a given residue that is accessible to water. These ASA values were then normalized by the maximum ASA value for a given amino acid to obtain RSA (Tien *et al.* 2013). Residues at sites with higher RSA have a larger part of the residue’s surface exposed to solvent and are generally closer to the protein’s surface while residues with lower RSA are closer to protein’s core. We defined sites with RSA ≤ 0.05 as buried sites and sites with RSA > 0.05 as exposed.

#### Packing density

We estimated residue packing density via the side-chain weighted contact number (WCN), defined as

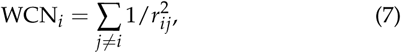

where *i* indicates the focal residue, *r*_*ij*_ is the distance between the geometric centers of the side chains of the focal residue *i* and of residue *j*, and the sum runs over all residues *j* in the protein. Since packing density tends to have a negative correlation with site entropy and RSA, here we use the inverse of WCN (iWCN = 1/WCN) for all correlation calculations, as was done previously by Shahmoradi *et al.* (2014).

We used side-chain WCN as defined above rather than WCN calculated from distances of C_*α*_ atoms because side-chain WCN provides the more robust determinant of evolutionary variation (Marcos and Echave 2015; Shahmoradi and Wilke 2016).

#### Score matching

To verify that our results were not caused by differences in stability between the designed and evolved sequences, we generated, for each template protein structure, subset alignments with matched stability scores between the designed and evolved sequences. We carried out this matching as follows. We first identified the intersection of the stability ranges between designed and evolved proteins. This intersection generally coincided with the stability range of the evolved proteins, i.e., evolved proteins had a narrower stability range than the corresponding designed proteins. We then identified, for each designed protein in that range, the evolved protein with the most similar score. Each designed protein was matched with exactly one unique evolved protein, and we stopped the matching step when there were no more designed or evolved proteins available for matching in the stability-range intersection. The resulting matched alignments consisted of between 19 and 433 sequences, with a median of 70.

### Data availability

All data and analysis scripts are available at: https://github.com/reductase4/evol_sim_vs_rosetta.git

## Results

To examine how evolutionary history affects the emergence of sequence variability, we analyzed two distinct sets of protein sequences, one produced by protein design (no evolutionary history, all amino acids are replaced at once) and one produced by evolutionary simulation (amino acids are introduced one at a time, and the protein need to remain viable at all times). Importantly, for both approaches we used the same methods to replace amino acids and evaluate the energy of the resulting protein structures, based on the fixed-backbone protein-design algorithm implemented in Rosetta (Kuhlman *et al.* 2003; Leaver-Fay *et al.* 2011). For each approach, we generated 500 sequences each from 38 template structures. To compare these sequences to natural sequences, we analyzed alignments of at least 50 homologs for each of the 38 protein structures, taken from Jackson *et al.* (2013) (see also Table S1). We found that the sequence divergence in the simulated alignments was comparable to that of the natural sequences, even though divergence in designed sequences was somewhat larger than divergence in evolved sequences (Figure 1).

**Figure 1.**
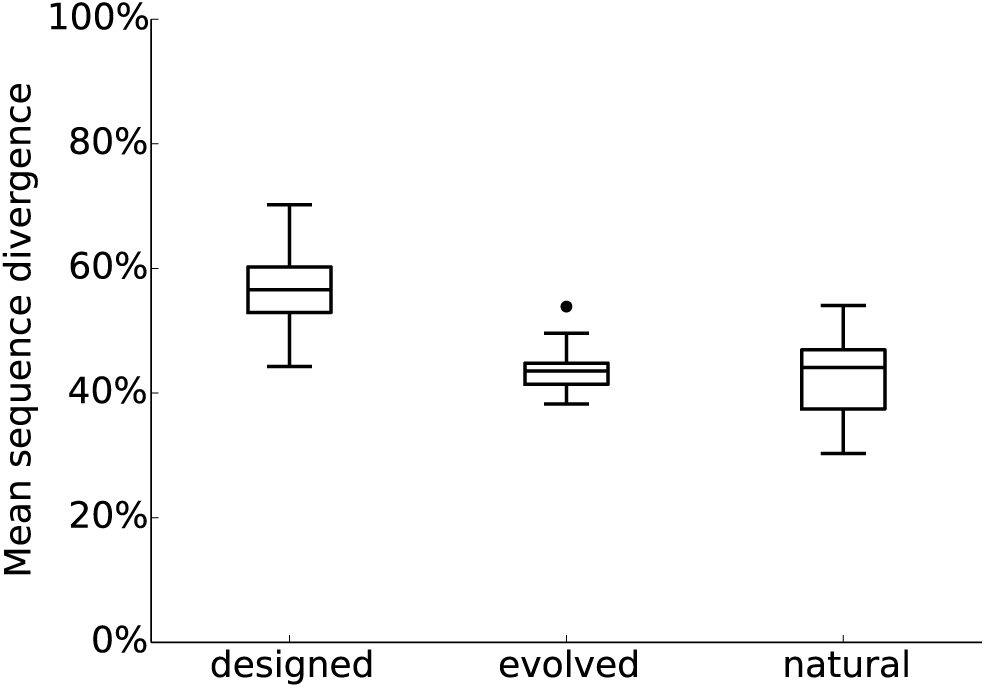
Mean sequence divergence from the starting sequence template in designed, evolved, and natural sequences. For natural sequences, we considered the sequence of the focal protein structure as the starting sequence template. The mean sequence divergence was higher in designed sequences than in both evolved (paired *t*-test, *p* < 2.2 × 10^−16^) and natural (paired *t*-test, *p* = 5.58 × 10^−13^) sequences, but it was comparable between evolved and natural sequences (paired *t*-test, *p* = 0.37).

### Amino-acid distributions

We first compared the overall amino-acid frequencies between natural and simulated sequences (Figure S1), because prior work comparing designed sequences to the same alignments of natural sequences had shown significant discrepancies, in particular for hydrophobic residues in buried sites (Jackson *et al.* 2013). We found that these discrepancies had mostly disappeared in our newly generated dataset. This difference may be due to an additional round of energy minimization we performed here, or (more likely) to recent updates to the energy function in Rosetta (Leaver-Fay *et al.* 2011).

However, there was now a substantial discrepancy for lysine, in that it was much more prevalent at exposed sites in designed sequences than in natural sequences. We also saw a moderate excess of arginines. We note that both are amino acids with a large number of rotamers, which the design algorithm biases towards. Importantly, there were only minor differences in amino-acid frequencies between evolved sequences and natural sequences. Because our evolutionary simulation made all mutations to different amino acids equally likely, we might have expected that amino acids encoded by four-or six-fold degenerate codon families would have been under-represented in comparison to their frequencies in natural alignments, which evolve at the DNA level and hence can be affected by codon degeneracy. Though these amino acids are not under-represented, neither in the evolutionary simulations nor in protein design, suggests that the energetic interactions between amino acids at different sites impose stronger constraints on amino-acid frequencies than does mutation bias. In summary, while evolutionary simulations outperformed protein design specifically for lysine and arginine, overall the differences between both approaches and natural sequences were minor.

While the analysis of aggregate amino-acid frequencies is useful as a first sanity check, it does not address whether either simulation approach places the correct amino acids at individual sites. Therefore, we next calculated the average Kullback-Leibler (KL) divergence for each protein, which quantifies the extent to which site-specific amino-acid distributions of simulated sequences are comparable to those of natural sequences. The lower the KL divergence, the more similar the distributions. As a control, we compared natural sequence to themselves, by randomly dividing each alignment into two groups and then comparing one to the other. We found that the evolutionary simulation produced sequences that were more similar to natural sequences than were the designed sequences (paired *t*-test, *p* < 2.2 ×10^−16^, Figure 2). Two exceptions (indicated as outliers in the middle boxplot of Figure 2) were chain D from the Sfi1p/Cdc31p complex (PDB ID: 2GV5) and initiation factor 5a (PDB ID:1XTD). This analysis suggests that accounting for evolutionary history is an important component in simulating realistic protein sequences.

**Figure 2.**
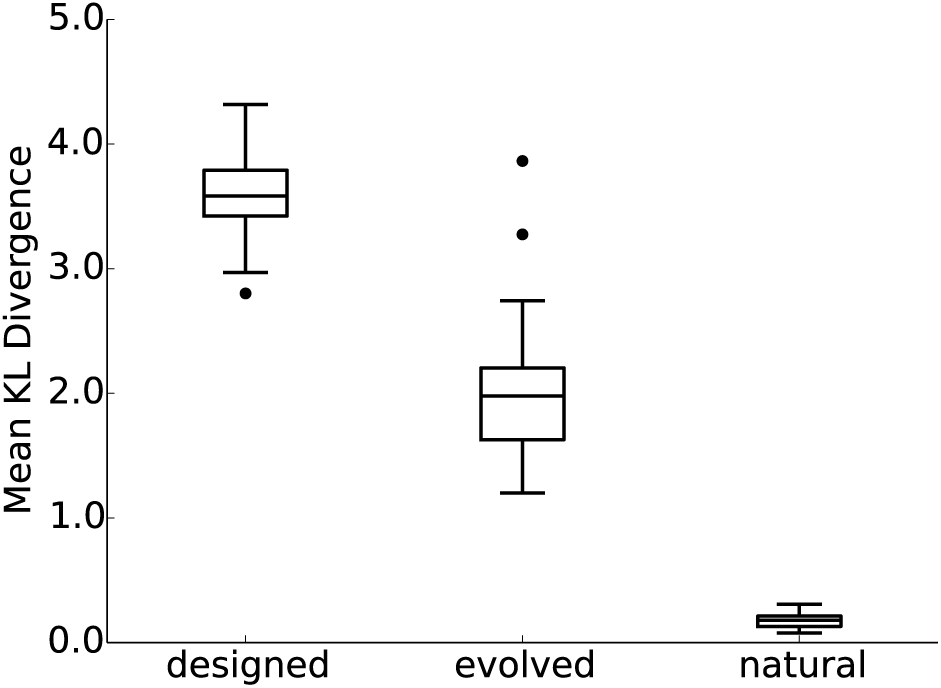
Mean Kullback-Leibler (KL) divergence of designed, evolved, and natural sequences to natural sequences. Natural sequences were compared to themselves by randomly splitting alignments into two groups and calculating the KL divergence between them. A lower KL divergence indicates that the amino acid distributions at individual sites are closer to that of natural proteins.

### Patterns of site variability

To investigate site-specific sequence variability we calculated the effective number of amino acids *n*_eff_ (Eq. 5) for each site in each protein. We found that sequences produced by simulating protein evolution showed similar overall variability to natural sequences (paired *t*-test, *p* = 0.68, Figure 3), whereas designed sequences had significantly less variability compared to natural sequences (paired *t*- test, *p* < 2.2 × 10^−16^, Figure 3). Importantly, designed sequences had a lower mean effective number of amino acids even though overall sequence divergence was higher in designed sequences than in evolved sequences (Figure 1). This seemingly paradoxical observation implies that designed sequences, relative to the evolved sequences, experienced changes at a larger number of sites but towards a smaller set of different amino acids.

**Figure 3.**
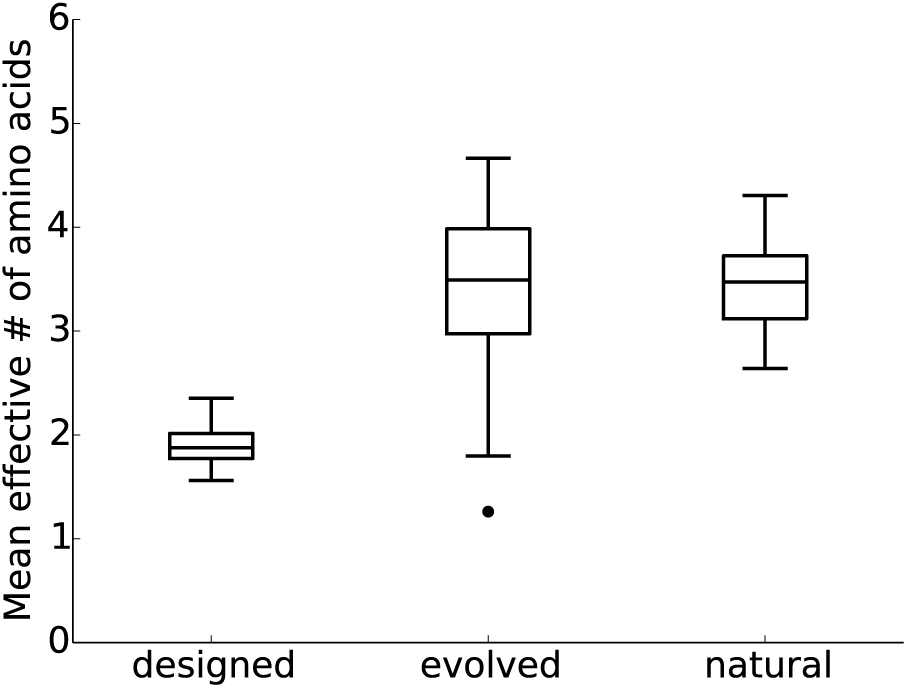
Mean effective number of amino acids for designed, evolved, and natural sequences. Evolved and natural sequences had comparable mean effective numbers of amino acids (paired *t*-test, *p* = 0.68) whereas designed sequences had significantly lower mean effective numbers (paired *t*-test, *p* < 2.2 × 10^−16^). One exception was chain D of the Sfi1p/Cdc31p complex (PDB ID: 2GV5, outlying data point in the boxplot for evolved sequences), for which evolutionary simulation yielded a much smaller mean effective number of amino acids relative to all other cases.

Just because two alignments have a comparable mean *n*_eff_ does not mean that the same sites are more variable or more conserved in the two alignments. Therefore, we next calculated the correlations between *n*_eff_ among alignments generated by different methods (designed sequences, evolved sequences, natural sequences). We found that the correlations in site variability between evolved and natural sequences were much higher than those between designed and natural sequences (Figure 4). The former were all positive and ranged between ~ 0.2 and 0.7, whereas the latter did not exceed ~ 0.3 and several fell below zero. Thus variable and conserved sites in evolved sequences tend to coincide with the same types of sites in natural sequence alignments, but the same is not true for designed sequences.

**Figure 4.**
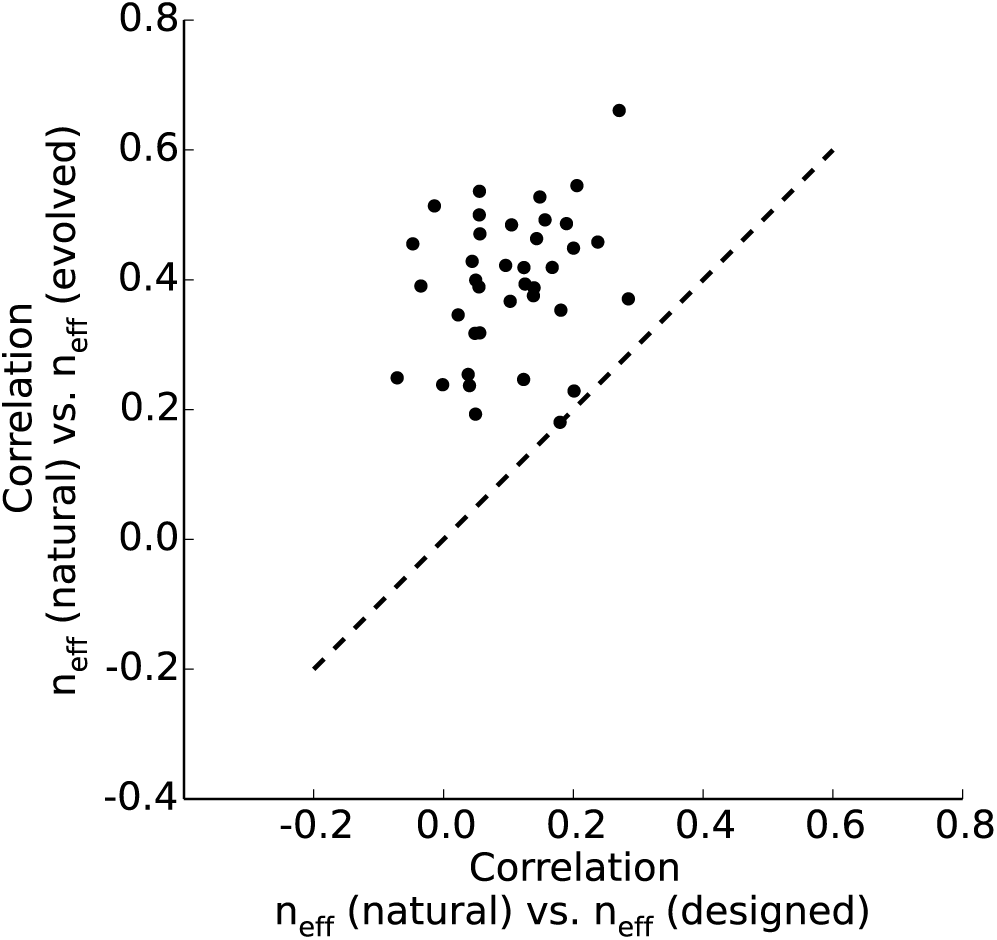
Spearman correlations between *n*_eff_ for natural and either evolved or designed sequences. Each dot represents the correlation coefficients for one protein. Correlations are substantially higher when comparing evolved to natural sequences than when comparing designed to natural sequences (paired *t* test, *p* < 2.2 × 10^−16^).

### Site variability in the context of structural features

Site-specific variability tends to correlate with features in the protein structure, most notably solvent exposure and packing density (Kimura and Ohta 1974; Yeh *et al.* 2013; Chang *et al.* 2013; Huang *et al.* 2014; Echave *et al.* 2016). Since these variability patterns are likely driven by biophysical constraints generated from the protein structure, we would expect that they would be recapitulated in computationally designed proteins. Yet, prior work had shown that site-variability patterns in designed proteins did not recapitulate the patterns observed in natural sequences— buried sites in designed sequences were not sufficiently conserved, or alternatively, exposed sites not sufficiently variable (Jackson *et al.* 2013).

We found here again that site variability in designed proteins did not appreciably correlate with relative solvent accessibility (RSA) (Figure 5). Correlations between *n*_eff_ and RSA ranged between −0.2 and 0.2. In comparison, for natural sequences these correlations fell mostly between 0.2 and 0.6. For evolved sequences, we observed even higher correlations, with most values falling between 0.5 and 0.7 (Figure 5). Results were similar when we used inverse Weighted Contact Number (iWCN) instead of RSA (Figure S2).

**Figure 5.**
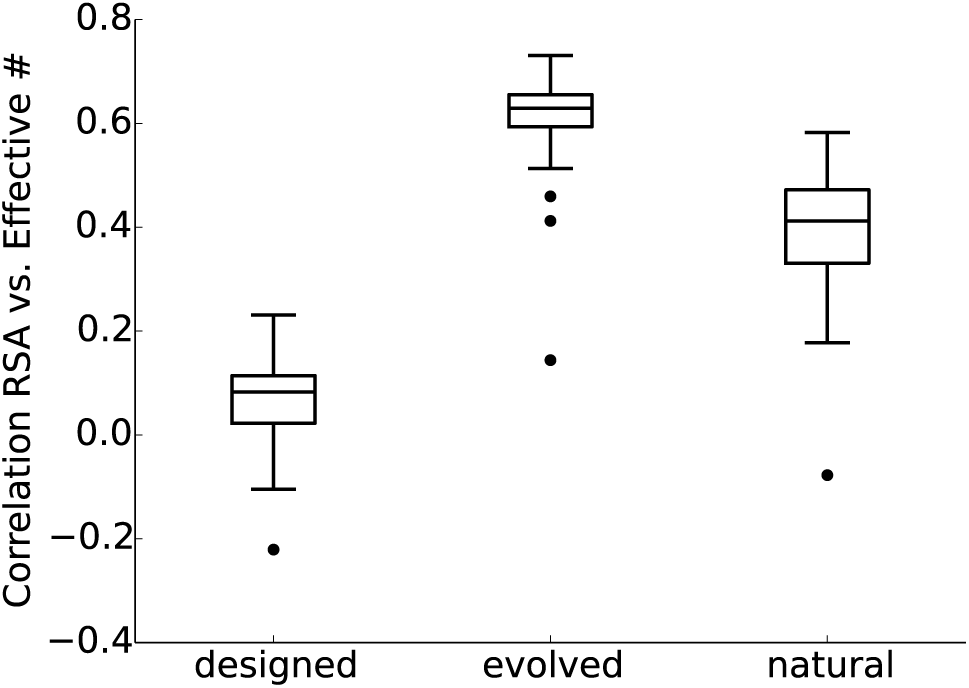
Spearman correlations between RSA and *n*_eff_ for designed, evolved, and natural sequences. Correlations for designed sequences were significantly lower than for natural sequences (paired *t*-test, *p* = 1.5 × 10^−14^). By contrast, correlations for evolved sequences were significantly *higher* than for natural sequences (paired *t*-test, *p* = 6.48 × 10^−12^). The individual correlation coefficients for each structure are shown in Figure S2.

One potential caveat to these findings is that designed and evolved sequences may fall into a different range of thermodynamic stability, assessed in our model via the Rosetta score. Indeed, even though we calibrated the stability threshold Δ*G*_thresh_ used during evolution to the mean stability for proteins designed to the same template (see Methods), we found that evolved proteins generally had a narrower range of stabilities than designed proteins and were, on average, more stable (Figure S3). In principle, these differences in stability distributions could be the cause for the other observed differences between designed and evolved sequences.

We addressed this caveat in two ways. First, we generated alignment subsets for each template structure such that each designed sequence retained in an alignment was matched one-to-one to a unique evolved sequence with comparable stability score and homologous structure. This procedure yielded near-identical stability distributions in most cases (Figure S4). Yet the observed pattern of RSA–*n*_eff_ correlations was virtually unchanged from that in the original dataset (Figure S5A). Second, for five arbitrarily chosen structures, we ran evolutionary simulations where we used the stability of the least stable designed structure (i.e., the maximum observed score among the designed structures) as stability threshold value. In those simulations, evolved structures were indeed much less stable than before (Figure S6). Yet again, there was virtually no change in the observed pattern of RSA–*n*_eff_ correlations (Figure S5B).

### Evolving designed sequences

To recap, we have found that by any metric considered, evolved sequence alignments looked much more realistic than designed sequence alignments. Further, we have seen that these differences between evolved and designed sequences are not caused by differences in protein stability. However, there do remain two potential reasons why we may have obtained these results: (i) evolutionary simulation adds an important element to sequence generation, one that is missed in the currently used protein design algorithm; (ii) evolved sequences look more similar to natural sequences simply because they have not diverged as much and thus retain much historical information about the natural ancestral sequence. To distinguish between these two scenarios, we ran an additional set of evolutionary simulations where we started each replicate evolutionary trajectory from one of the previously designed sequences. We performed this additional set of simulations for a subset of 10 arbitrarily chosen structures (Table S1). We refer to sequences generated in this manner as “evolved from design.” Importantly, the mean sequence divergence in these sequences was significantly higher than in the corresponding natural or evolved sequences, and almost as high as in the designed sequences (Figure S7).

From these additional simulations, we found that the evolutionary process produced more naturally-looking alignments even when designed proteins were used as starting points (Figure 6). Sequences that were evolved from design had a lower KL divergence than designed sequences (Figure 6A), a higher mean effective number of amino acids (Figure 6B), higher correlations of site-variability patterns to natural alignments (Figure 6C), and higher correlations between *n*_eff_ and RSA (Figure 6D). However, by all these metrics, sequences evolved from design were intermediate between designed sequences and sequences evolved from a natural sequence. These intermediate metrics reflect that the designed sequences have a site-wise preference for a smaller set of amino acids than do the evolved sequences (Figure 3), and this preference is only partly undone by subsequent evolution. While evolution could expand the set of acceptable amino acids at some sites, other sites remained frozen in the narrow area of sequences space supplied by design and might require much longer evolutionary simulations to become unfrozen.

**Figure 6.**
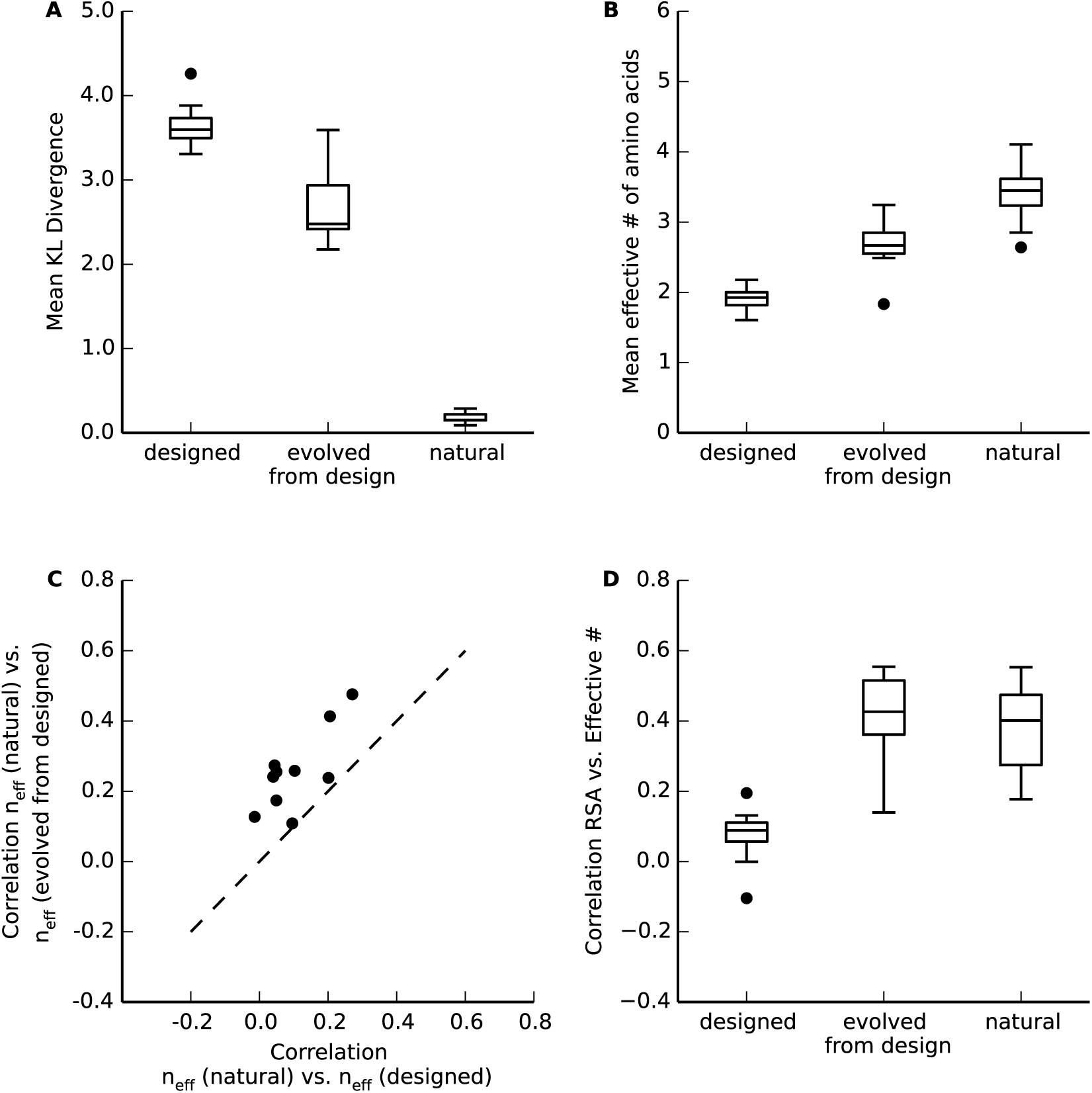
Comparison between designed sequences and sequences evolved from design. (A) Mean Kullback-Leibler (KL) divergence of designed, evolved from design, and natural sequences. The KL divergence was lower for evolved from design sequences than for designed sequences (paired *t*-test, *p* = 2.57 × 10^−6^). (B) Mean effective number of amino acids for designed, evolved from design, and natural sequences. The mean effective number *n*_eff_ was higher for evolved from design sequences than for designed sequences (paired *t*-test, *p* = 1.91 × 10^−5^) (C) Spearman correlations of *n*_eff_ between natural and evolved from design sequences and natural and designed sequences. Each dot represents one correlation coefficient value for one protein. Correlations with *n*_eff_ from natural sequences were significantly higher for *n*_eff_ from evolved from design sequences than for *n*_eff_ from designed sequences (paired *t*-test, *p* = 6.23 × 10^−5^). (D) Spearman correlations between site entropy and RSA of designed, evolved from design, and natural sequences. Correlations were significantly higher for evolved from design sequences than designed sequences (paired *t*-test, *p* = 1.7 × 10^−7^), however correlations for evolved from design and natural sequences were comparable (paired *t*-test, *p* = 0.62).

### Using simulated sequences as a predictor of site variability

Finally, we asked how well *n*_eff_ from simulated sequences performed as predictor of natural site variability relative to the other commonly used predictors RSA and WCN. In agreement with prior work (Shahmoradi *et al.* 2014; Jackson *et al.* 2016), we saw that the *n*_eff_ from designed sequences performed poorly relative to RSA (Figure 7A). However, the *n*_eff_ from evolved sequences performed similarly to RSA (Figure 7B), and *n*_eff_ from evolved from design sequences displayed intermediate performance (Figure 7C). Results were similar for the inverse weighted contact number (iWCN), except that correlations between *n*_eff_ and iWCN tended to be somewhat stronger than they were between *n*_eff_ and RSA (Figure 7D, E, F), consistent with a large body of prior work finding that WCN tends to correlate more strongly with evolutionary variation than RSA does (Yeh *et al.* 2013, 2014; Shahmoradi *et al.* 2014; Marcos and Echave 2015; Echave *et al.* 2016). Unlike RSA, WCN measures both the local structural constraint around a residue and the global arrangement of amino acids in the entire structure Shahmoradi and Wilke (2016). The elevated correlations seen with WCN relative to RSA may thus in part be driven by factors other than folding constraints, such as the location of active sites in enzymes (Jack *et al.* 2016).

**Figure 7.**
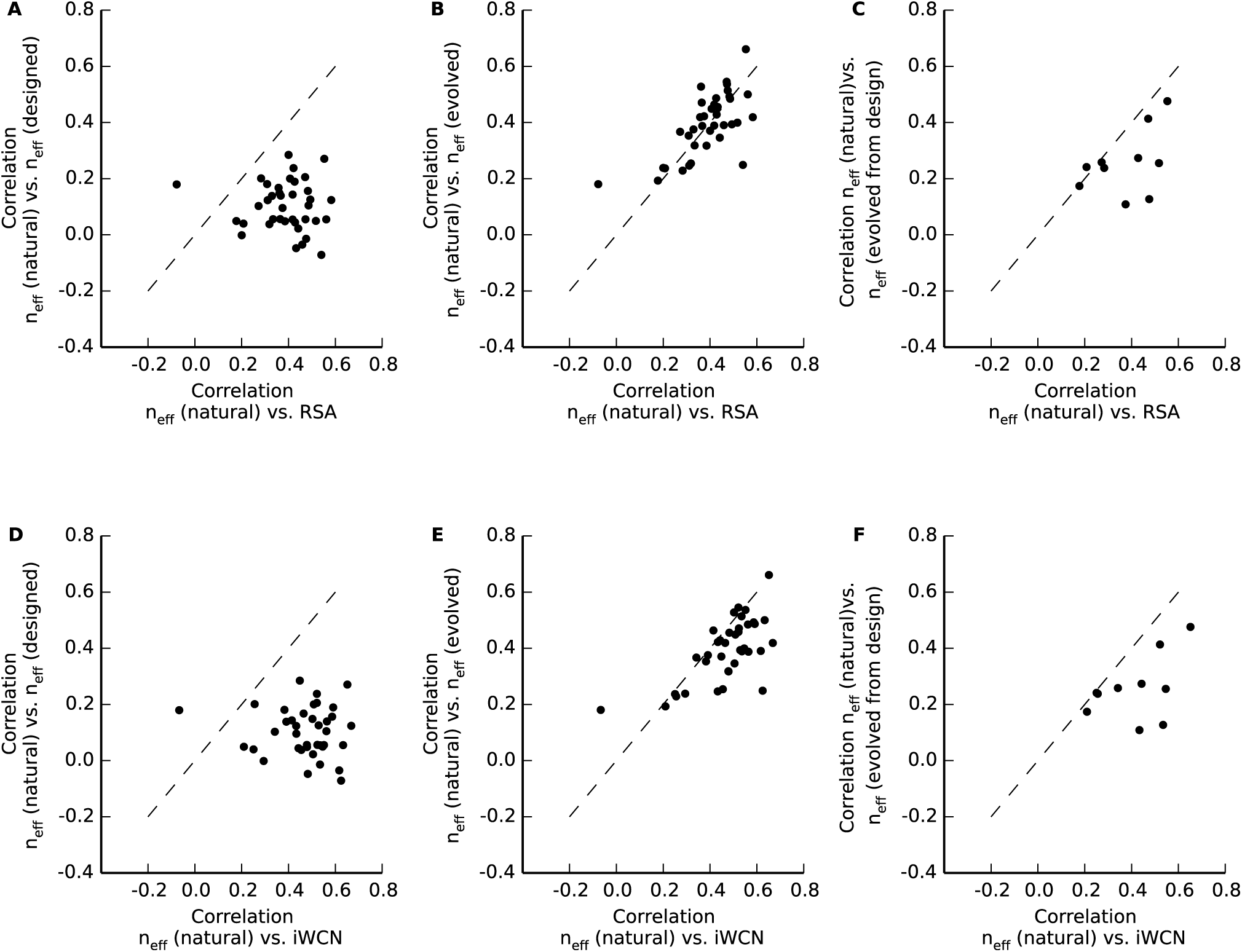
Spearman correlations between *n*_eff_ for natural sequences and *n*_eff_ for simulated sequences values versus Spearman correlations between *n*_eff_ for natural sequences structural measures RSA (A), (B), (C) or iWCN (D), (E), (F). These comparisons allow us to asses whether *n*_eff_ in natural sequences correlates better with *n*_eff_ in simulated sequences or with structural measures. We find that evolved sequences perform similarly to RSA and iWCN, but designed sequences do not.

## Discussion

We have examined how evolutionary history affects the emergence of sequence variability, by comparing protein alignments generated by two different methods. The first was a traditional protein-design algorithm implemented in RosettaDesign (Kuhlman *et al.* 2003). The other was an evolutionary method that used the Rosetta energy function to simulate protein evolution according to population-genetics principles (Teufel and Wilke 2017). Both methods impose the same thermodynamic constraints on the simulated sequences but they impose different constraints in their sampling of sequence space. For both methods, we compared the simulated alignments to homologous alignments of natural sequences. We found that sequences generated by simulated evolution displayed site-specific variation quite similar to that of natural sequences, whereas patterns of variation were substantially different in designed sequences. Our evolved sequences also showed correlations between sequence variation and residue burial or packing similar to that of natural sequences whereas the designed sequences did not. Finally, by simulating additional evolution with designed sequences as starting points, we demonstrated that the improvements in site-variability metrics were caused by the evolutionary process itself, and not by insufficient divergence from the starting sequence during the evolutionary simulations.

While both evolutionary and protein-design methods can be used to generate proteins that fold and function, they differ in how they explore sequence space. In particular, the widely used RosettaDesign (Kuhlman *et al.* 2003) takes a template structure, strips all residue side chains, and then simultaneously replaces them by different side chains. Subsequently additional side-chain replacements are made with the goal to maximize protein stability. This process of designing a sequence can be repeated a number of times to generate a set of diverse sequences. This and similar approaches have proven fruitful for protein engineering. For example, protein design methods have been used to engineer proteins that bind an influenza virus (Fleishman *et al.* 2011), to create enzymes (Röthlisberger *et al.* 2008), and to develop novel protein folds (Kuhlman *et al.* 2003). However, this method of generating functional proteins is very unlike how natural systems generate them. Natural proteins accumulate changes mostly sequentially, one mutation at a time, rather than having their entire sequence modified in a single instance.

Protein design is frequently billed as being able to explore a much larger sequence space than does evolution, because the design process starts from scratch with a random sequence and thus in theory should be able to reach any place in sequence space whereas evolution can only explore near neighbors to already existing sequences (Huang *et al.* 2016). However, we did not find this to be the case in our simulations. Specifically, even though our designed sequences had a higher mean sequence divergence than did evolved or natural sequences (Figure 1), designed sequences had, on average, smaller effective numbers of amino acids at each site (Figure 3). Thus, on average, individual sites were less variable under design than they were under evolution, natural or simulated. We were able to explain this unexpected result by investigating how exactly the initial sequences are chosen under design. The initial sampling of side chains is done from a rotamer library (Kaufmann *et al.* 2010), rather than uniformly from the 20 amino acids. Consequently, amino acids with many rotamers are over-represented in the initially chosen sequence. The subsequent optimization algorithm is constrained by the biased pool of initial sequences and cannot undo the unequal sampling. We can see the remnants of the oversampling of certain amino acids in the excess of lysine and arginine in the overall amino-acid frequencies (Figure S1), as both are amino acids with a very large number of rotamers. When we subjected the designed sequences to further evolution a wider range of substitutions became allowed. Therefore, we saw the evolved-from-design sequences approaching site variabilities similar to those displayed by the original evolved and natural sequences (Figure 6).

Our findings are consistent with recent works on evolutionary entrenchment (Pollock *et al.* 2012; McCandlish *et al.* 2016, 2015; Shah *et al.* 2015; Goldstein and Pollock 2016), which argue that the propensity of a protein to acquire position-specific substitutions varies over time, as previously accumulated mutations become entrenched in the protein structure and slowly alter the constraints imposed on other amino acids in the structure. Here, entrenchment was visible in particular for designed and evolved-from-design sequences, which were highly and somewhat biased by the initial sampling process of protein design.

Even though our evolutionary simulations considered both structural constraints and a realistic, sequential sampling of sequence space, we found that they did not fully capture the relationship between sequence variability, buriedness, and packing density observed in natural sequences. In particular, site-specific variability was correlated more strongly with RSA in our evolved sequences than is observed in nature. This exaggerated relationship between sequence variability and RSA may be caused by our model’s simplistic assumption that selection acts exclusively on protein stability. In natural organisms, numerous selection pressures beyond just protein stability act on protein sequences (Thorne 2007; Teufel *et al.* 2012; Serohijos and Shakhnovich 2013, 2014; Chi and Liberles 2016; Echave and Wilke 2017), and these other selection pressures should weaken the observed relationship between RSA and sequence variability.

There are several caveats to our work. First, throughout this project, we have used a fixed backbone model, even though allowing for a flexible backbone during design may produce more natural-looking sequences (Jackson *et al.* 2013; Ollikainen and Kortemme 2013). However, since we used the same fixed backbone model for both protein design and protein evolution, we don’t expect it to have much bearing on the observed differences between designed and evolved sequences. Moreover, Jackson *et al.* (2013) had previously shown that even with flexible backbone design the correlation between solvent accessibility and site variability was lower than observed in natural sequences. Second, the accelerated origin-fixation model we used for evolution changes the order in which substitutions are accumulated, but we expect this reordering to be of little consequence for the metrics considered in this work (Teufel and Wilke 2017). Third, the amount of divergence generated during simulated evolution depends on the length of time for which the simulations are run, and thus is highly dependent on the parameter choices made for the simulations. We chose to run the evolutionary simulations for 5000 substitutions in each trajectory (for proteins of at most several hundred amino acids in length), to ensure that the amount of divergence generated during simulation would exceed that observed in typical natural homologs. Finally, it is possible that a modified design algorithm that samples initial sequences from a uniform distribution of amino acids rather than from a distribution of rotamers would produce sequence alignments on par with those we generated here under simulated evolution. Testing this hypothesis, however, falls beyond the scope of the present paper.

## Acknowledgments

This work was supported by National Institutes of Health Grants R01 GM088344 and National Science Foundation Cooperative agreement no. DBI-0939454 (BEACON Center). Computational resources were provided by The University of Texas at Austin’s Center for Computational Biology and Bio-informatics and the Stampede cluster at the Texas Advanced Computing Center. Q.J. was a visiting scholar at UT Austin (September 2015–September 2017) sponsored by the China Scholarship Council (CSC). E.L.J. was supported by NSF GRFP DGE-1110007.

---

**Table S1.**
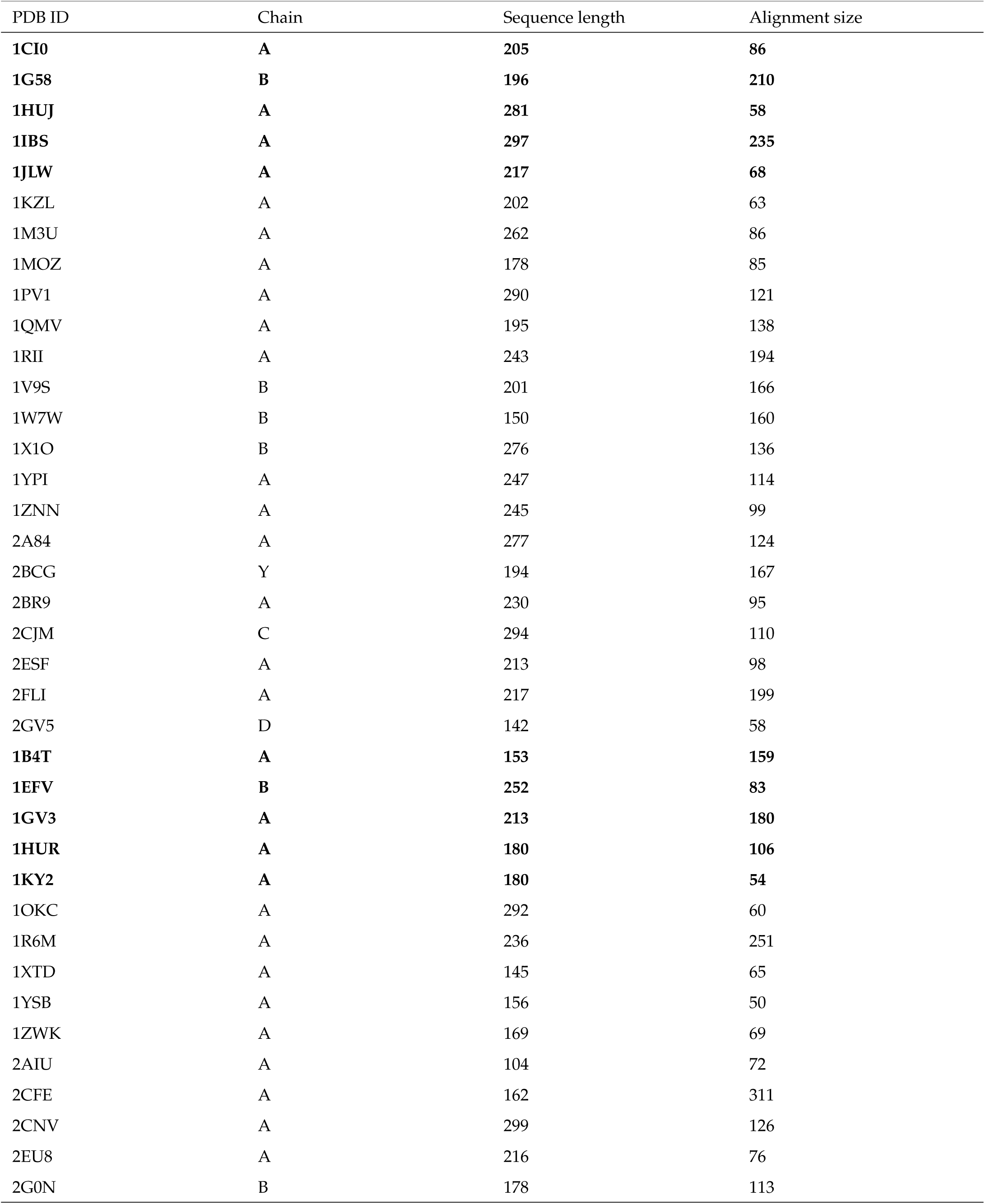
PDB structures used in this study. For each structure, we show the PDB identifier (“PDB ID”), the chain, the length of the corresponding sequence in amino acids, and the number of sequences in the alignment of homologous natural sequences (“alignment size”). Structures that were subjected to the additional “evolved from design” protocol are highlighted in bold.

| PDB ID | Chain | Sequence length | Alignment size |
| --- | --- | --- | --- |
| **1CI0** | **A** | **205** | **86** |
| **1G58** | **B** | **196** | **210** |
| **1HUJ** | **A** | **281** | **58** |
| **1IBS** | **A** | **297** | **235** |
| **1JLW** | **A** | **217** | **68** |
| 1KZL | A | 202 | 63 |
| 1M3U | A | 262 | 86 |
| 1MOZ | A | 178 | 85 |
| 1PV1 | A | 290 | 121 |
| 1QMV | A | 195 | 138 |
| 1RII | A | 243 | 194 |
| 1V9S | B | 201 | 166 |
| 1W7W | B | 150 | 160 |
| 1X1O | B | 276 | 136 |
| 1YPI | A | 247 | 114 |
| 1ZNN | A | 245 | 99 |
| 2A84 | A | 277 | 124 |
| 2BCG | Y | 194 | 167 |
| 2BR9 | A | 230 | 95 |
| 2CJM | C | 294 | 110 |
| 2ESF | A | 213 | 98 |
| 2FLI | A | 217 | 199 |
| 2GV5 | D | 142 | 58 |
| **1B4T** | **A** | **153** | **159** |
| **1EFV** | **B** | **252** | **83** |
| **1GV3** | **A** | **213** | **180** |
| **1HUR** | **A** | **180** | **106** |
| **1KY2** | **A** | **180** | **54** |
| 1OKC | A | 292 | 60 |
| 1R6M | A | 236 | 251 |
| 1XTD | A | 145 | 65 |
| 1YSB | A | 156 | 50 |
| 1ZWK | A | 169 | 69 |
| 2AIU | A | 104 | 72 |
| 2CFE | A | 162 | 311 |
| 2CNV | A | 299 | 126 |
| 2EU8 | A | 216 | 76 |
| 2G0N | B | 178 | 113 |

**Figure S1.**
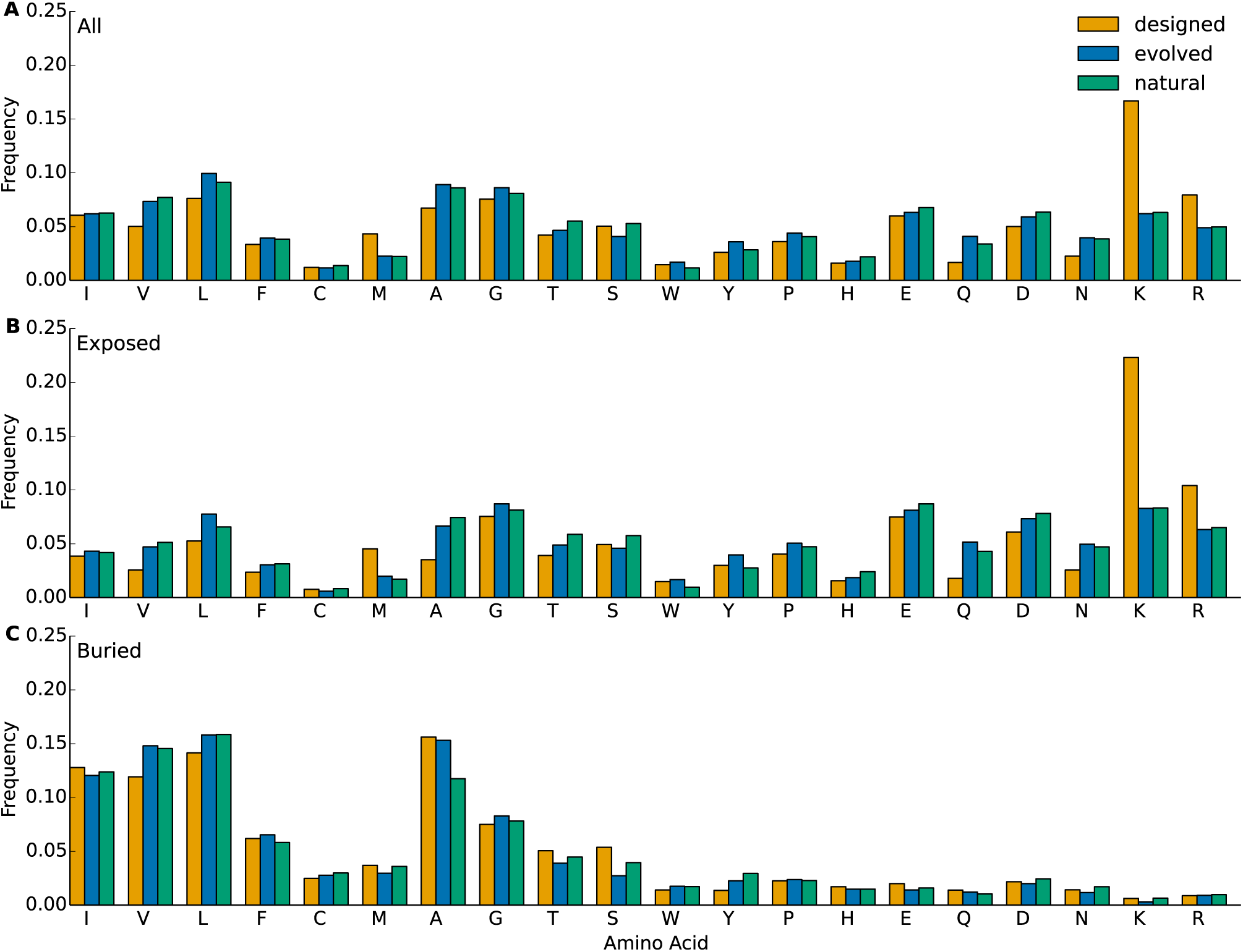
Amino-acid frequencies in designed, evolved, and natural sequences. (A) Aggregate frequencies of residues at all sites in all alignments. (B) Aggregate frequencies of residues at exposed sites (defined as sites with RSA > 0.05) in all alignments. (C) Aggregate frequencies of residues at buried sites (defined as sites with RSA ≤ 0.05) in all alignments.

**Figure S2.**
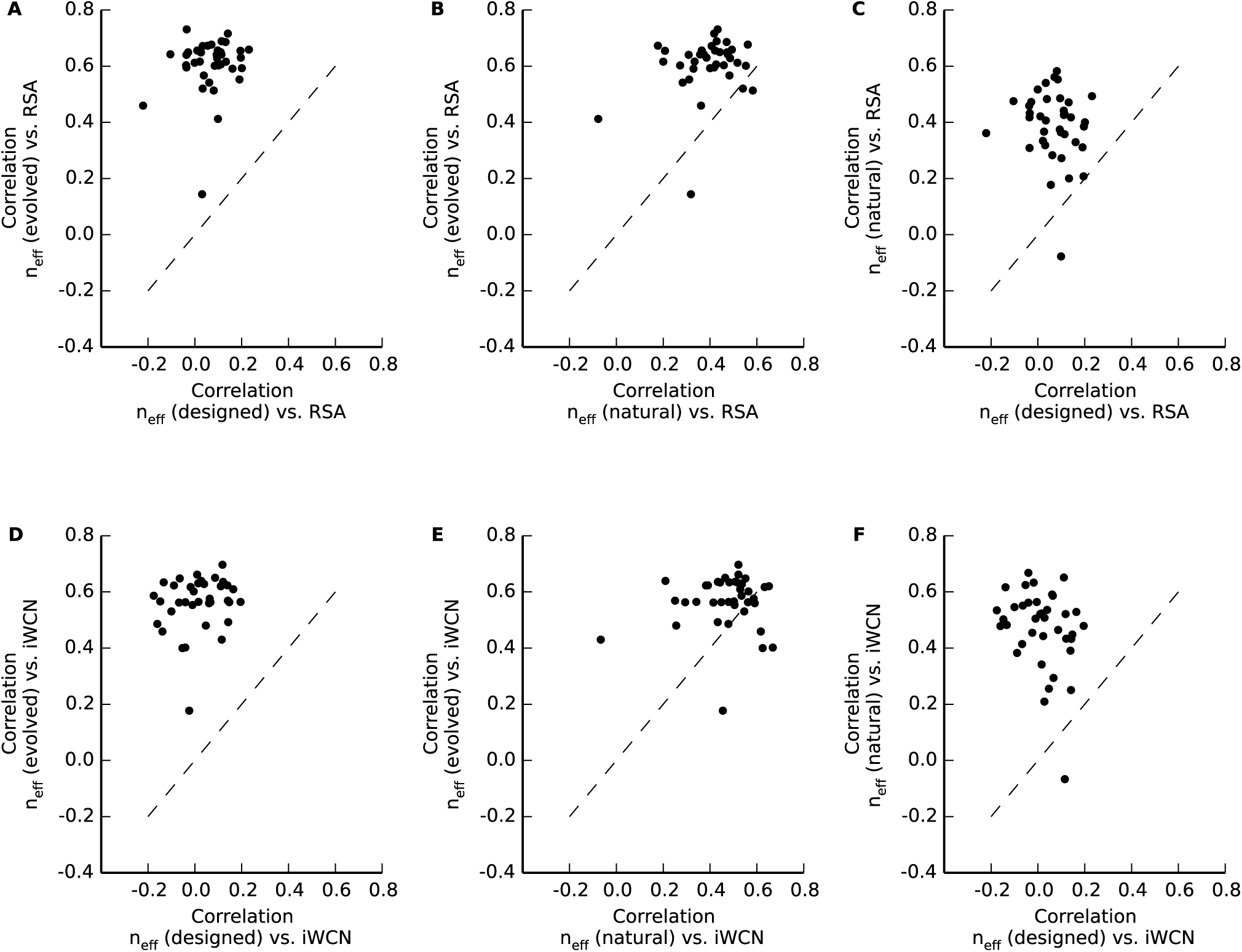
Spearman correlations between site entropy and RSA or iWCN. (A), (B), (C) Spearman correlations between site entropy and RSA. (D), (E), (F) Spearman correlations between site entropy and iWCN. Each dot represents one pair of correlation coefficients for one protein.

**Figure S3.**
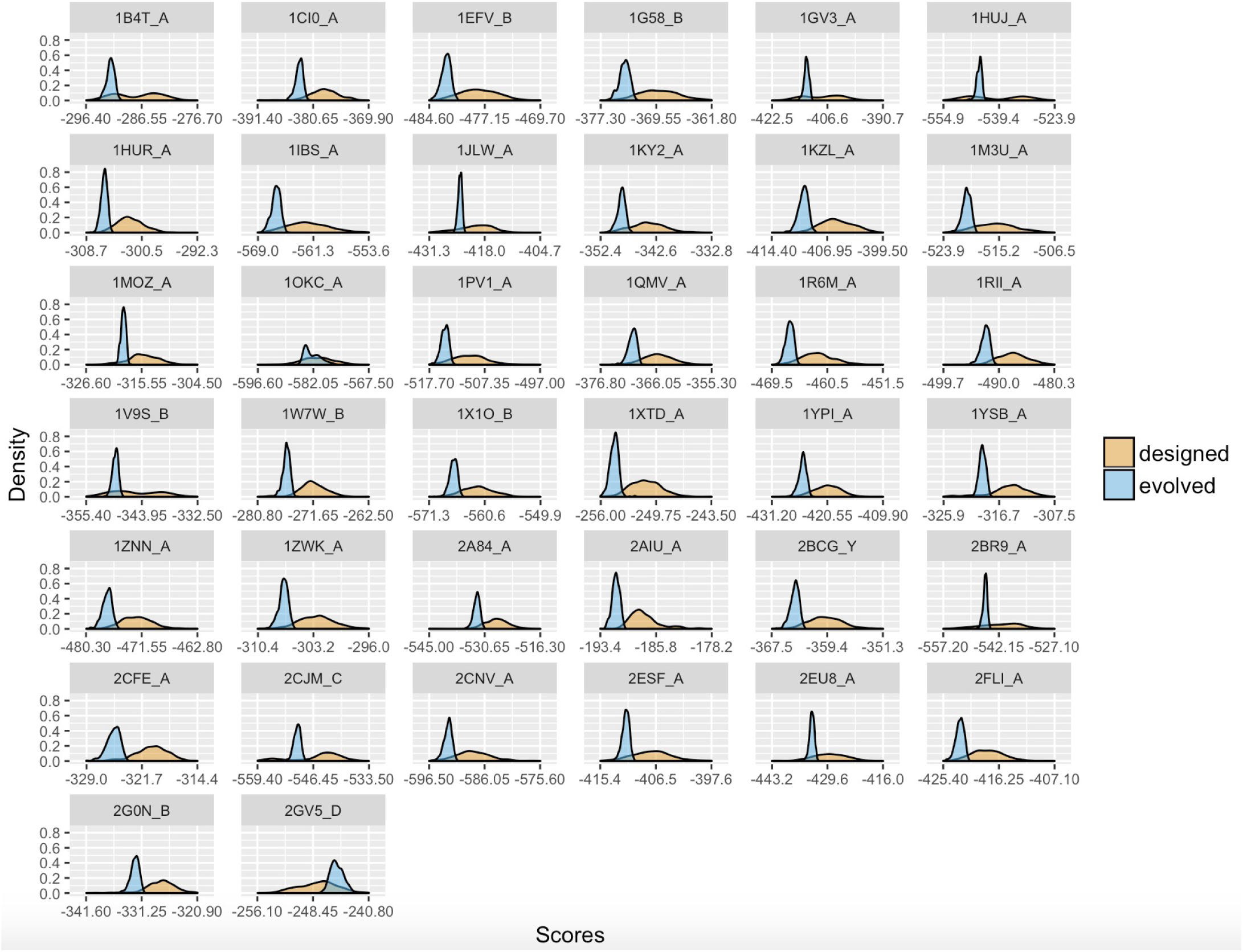
Density distributions of scores calculated from designed and evolved structures. The evolved structures were evolved using the mean score from the designed structures as the threshold value *G*_thresh_. On average, evolved structures occupied a narrower range of stability scores and were more stable than designed structures.

**Figure S4.**
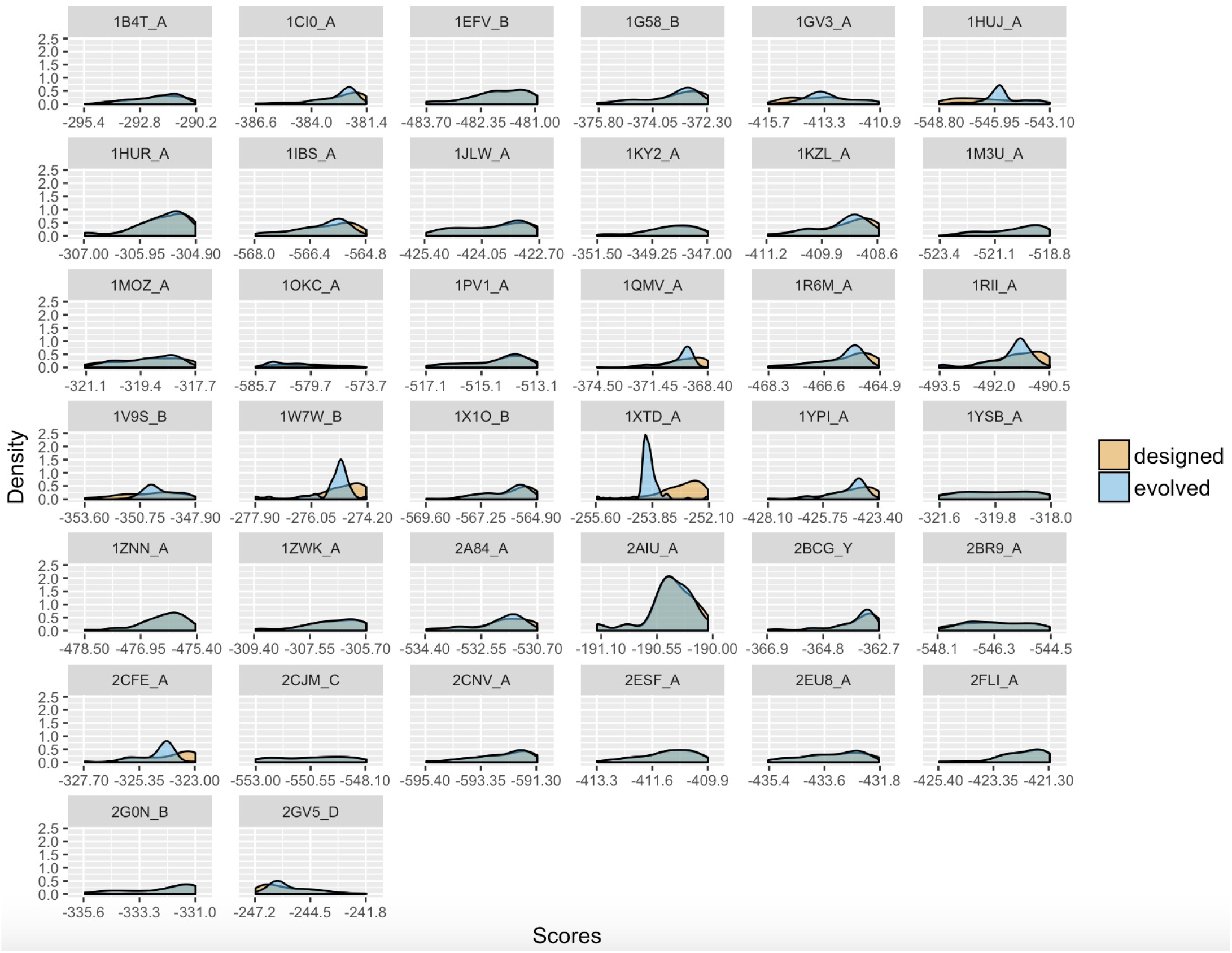
Density distributions of scores calculated from designed and evolved structures after a subset of designed structures was matched to evolved structures with most similar scores. In most cases, the matching produced distributions that were very similar or nearly identical. Notable exceptions were 1RII_A, 1W7W_B, 1XTD_A, and 2CFE_A.

**Figure S5.**
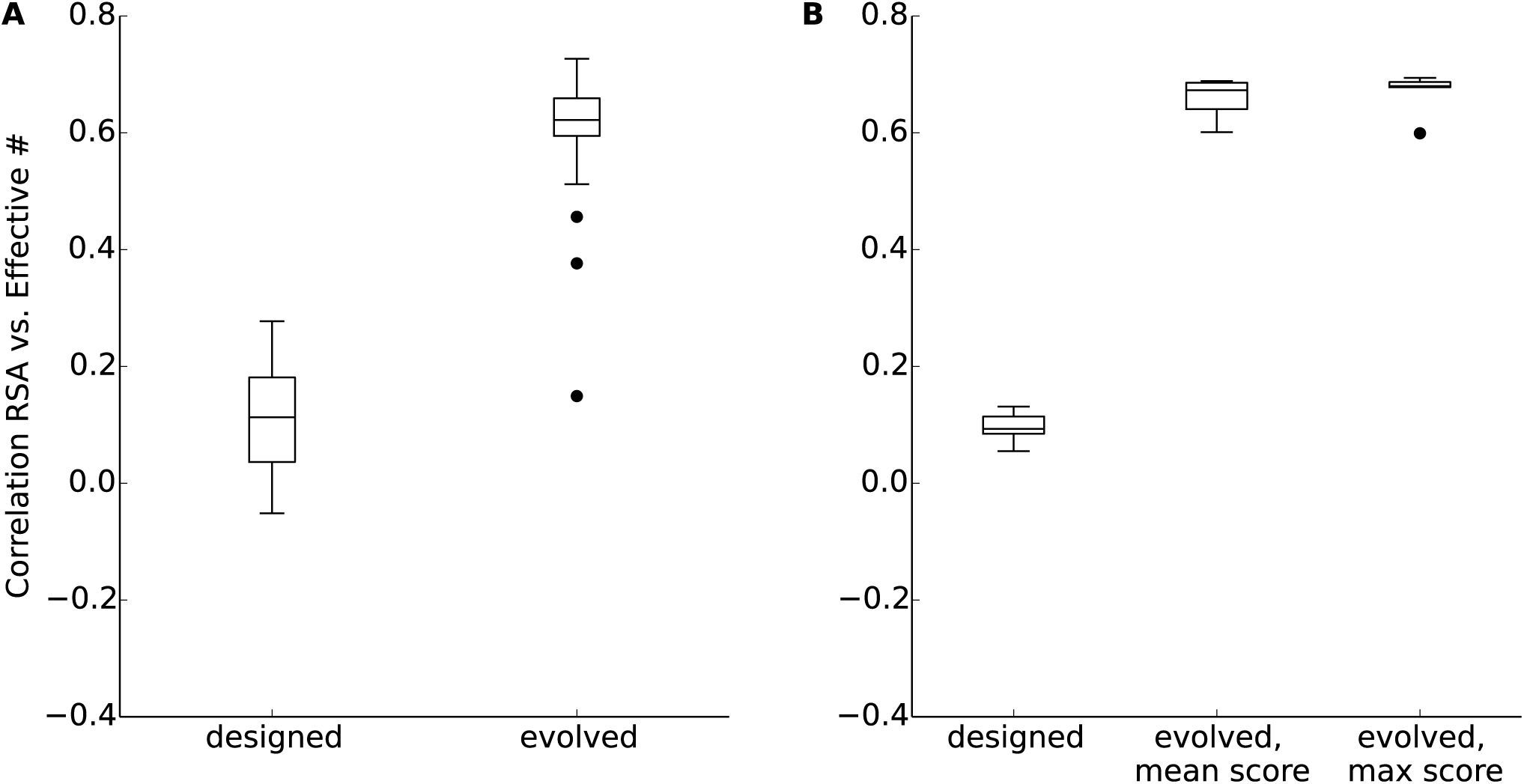
Spearman correlations between RSA and *n*_eff_. (A) Spearman correlations between RSA and *n*_eff_ for proteins with matched score distributions. See Fig. S3 for score distributions before matching and Fig. S4 for score distributions after matching. Evolved sequences had significantly higher correlations than designed sequences (paired *t*-test, *p* = 2.2 × 10^−16^). (B) Spearman correlations between RSA and *n*_eff_ for five proteins (1B4T_A, 1EFV_B, 1GV3_A, 1HUR_A, 1KY2_A) for which we reran the evolutionary simulations using the maximum (least stable) score from protein design as the threshold value *G*_thresh_. Shown are the distributions of correlation coefficients for the designed proteins, the proteins evolved using the mean score as threshold value, and the proteins evolved using the maximum score as threshold value. The score distributions from the latter simulations are shown in Fig. S6. Sequences evolved with either threshold value had significantly higher correlations than designed sequences (paired *t*-test, *p* = 2.34 × 10^−6^ and *p* = 3.4 × 10^−6^, respectively). There was no detectable difference in correlation strengths between the two datasets of evolved sequences (paired *t*-test, *p* = 0.36).

**Figure S6.**
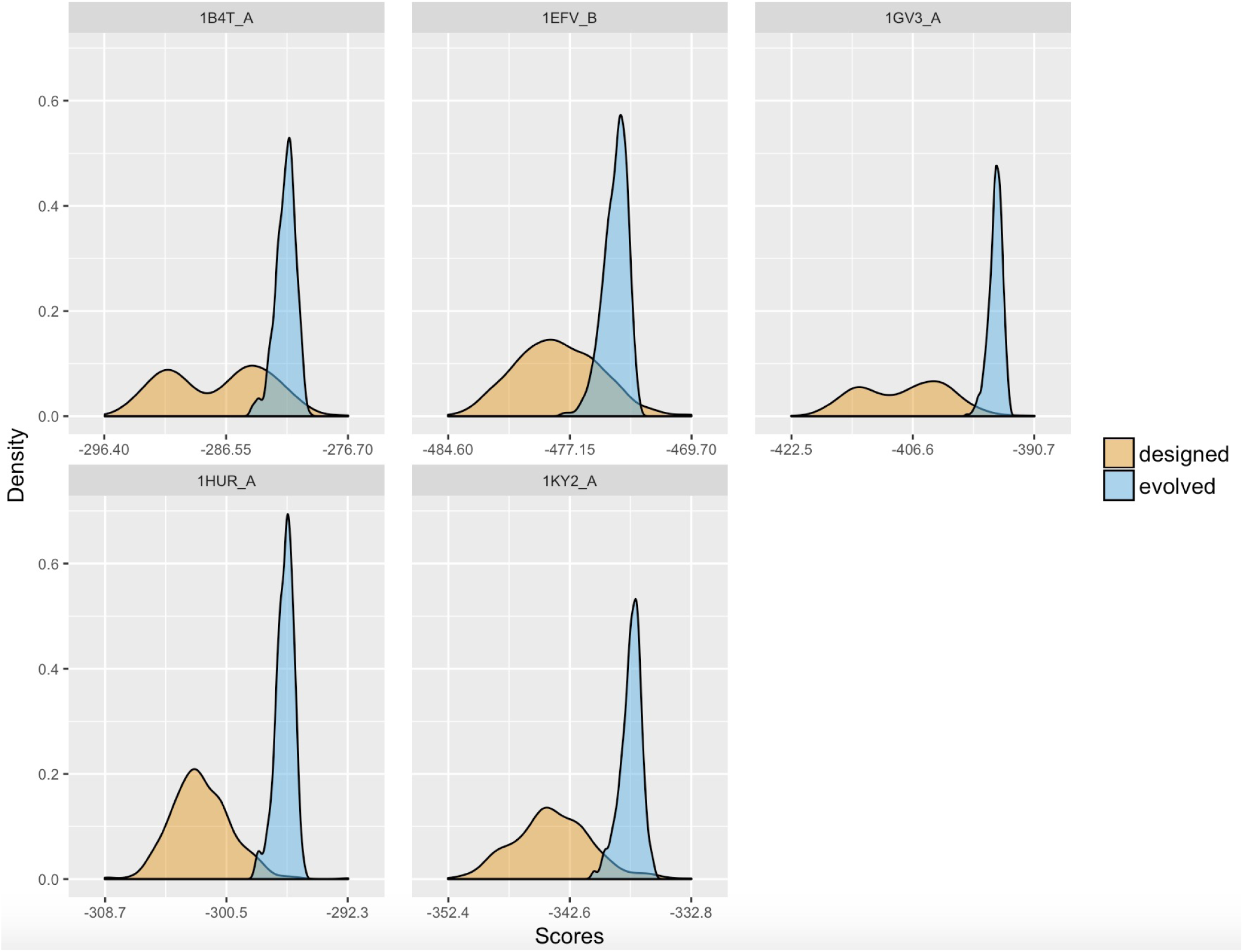
Density distribution of scores calculated from designed and evolved structures. The evolved structures were evolved with the maximum score from the designed structures used as the threshold value *G*_thresh_.

**Figure S7.**
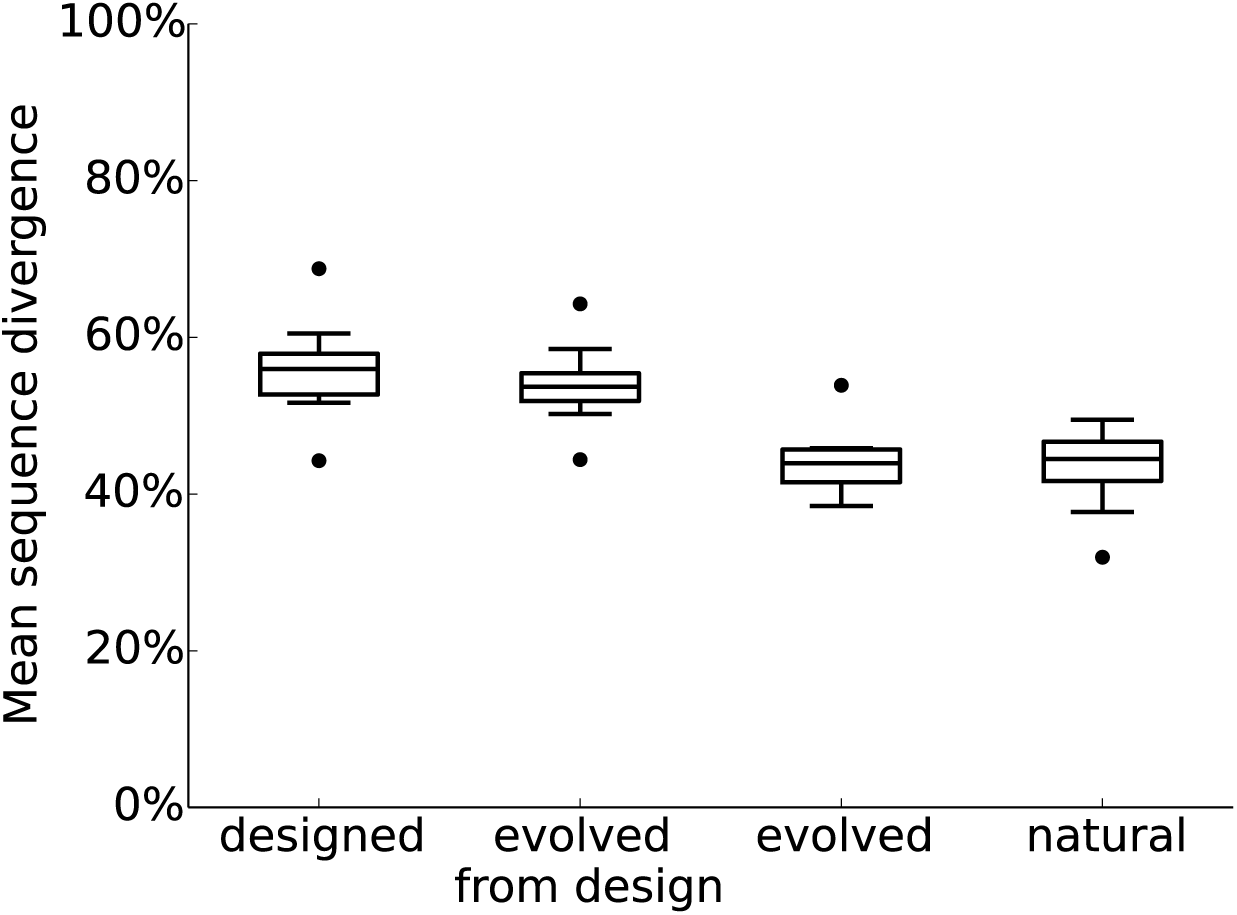
Mean sequence divergence from the starting sequence template in designed, evolved from designed, evolved, and natural sequences. Results are shown only for the 10 structures subjected to the additional “evolved from design” protocol (see Table S1). Designed and evolved-from-design sequences had significantly higher mean sequence divergence than natural sequences (*p* = 0.00014 and *p* = 0.00018, respectively, paired *t*-tests). However, designed sequences also had significantly higher mean sequence divergence than evolved-from-design sequences (paired *t*-test, *p* = 0.0019).

